# NLR signaling network mediates immunity to diverse plant pathogens

**DOI:** 10.1101/090449

**Authors:** Chih-Hang Wu, Ahmed Abd-El-Haliem, Tolga O. Bozkurt, Khaoula Belhaj, Ryohei Terauchi, Jack H. Vossen, Sophien Kamoun

## Abstract

Plant and animal nucleotide-binding domain and leucine-rich repeat-containing (NLR) proteins often function in pairs to mediate innate immunity to pathogens. However, the degree to which NLR proteins form signaling networks beyond genetically linked pairs is poorly understood. In this study, we discovered that a large NLR immune signaling network with a complex genetic architecture confers immunity to oomycetes, bacteria, viruses, nematodes, and insects. The network emerged over 100 million years ago from a linked NLR pair that diversified into up to one half of the NLR of asterid plants. We propose that this NLR network increases robustness of immune signaling to counteract rapidly evolving plant pathogens.

---

Plants and animals rely on nucleotide-binding domain and leucine-rich repeat-containing (NLR) proteins to activate immune responses to invading pathogens (1-3). NLR are among the most diverse and rapidly evolving protein families in plants (4, 5). They are modular proteins that broadly fall into two classes based on their N-terminal domain, which is either a Toll-interleukin 1 receptor (TIR) or a coiled coil (CC) domain (6). Most plant disease resistance genes encode NLR receptors that detect effector proteins secreted by pathogens either by directly binding them or indirectly via effector-targeted host proteins (3, 7). An emerging model is that “sensor” NLR dedicated to detecting pathogen effectors require “helper” NLR to initiate immune signaling resulting in a hypersensitive cell death response that restricts pathogen invasion (8-12). Although paired NLR have been described across flowering plants, the degree to which plant NLR have evolved to form higher order networks is poorly known.

The Solanaceae form one of the most species-rich plant families that includes major agricultural crops, such as potato, tomato and pepper (13). Solanaceae genomes harbor hundreds of NLR-type genes, over 20 of which have been demonstrated to confer resistance to infection by diverse and destructive pathogens and pests, including the Irish potato famine agent *Phytophthora infestans* (14, 15). As part of a study performed in *Nicotiana benthamiana* to identify genetic components required for resistance to *P. infestans* conferred by the potato NLR-type gene *Rpiblb2* (16, 17), we discovered that another NLR protein, NRC4 (<u>N</u>LR <u>r</u>equired for <u>c</u>ell death <u>4</u>) (9, 18), is required for Rpi-blb2 function (Fig. 1). Silencing of *NRC4* compromised Rpi-blb2 resistance to *P. infestans* (Fig. 1A) and hypersensitive cell death to the *P. infestans* AVRblb2 effector (Fig. 1B) (17). This phenotype was rescued by a silencing-resilient synthetic *NRC4* gene (Fig. 1C-D, Fig. S1A-B). *NRC4*-silencing did not affect Rpi-blb2 accumulation (Fig. S1C). Mutation in the ATP-binding p-loop motif of both Rpi-blb2 and NRC4 abolished their activities (Fig. S2). Thus a strict sensor/helper model where only one NLR requires ATP binding (19, 20) is too simple to explain the interaction between the Rpi-blb2/NRC4 pair.

**Fig. 1.**
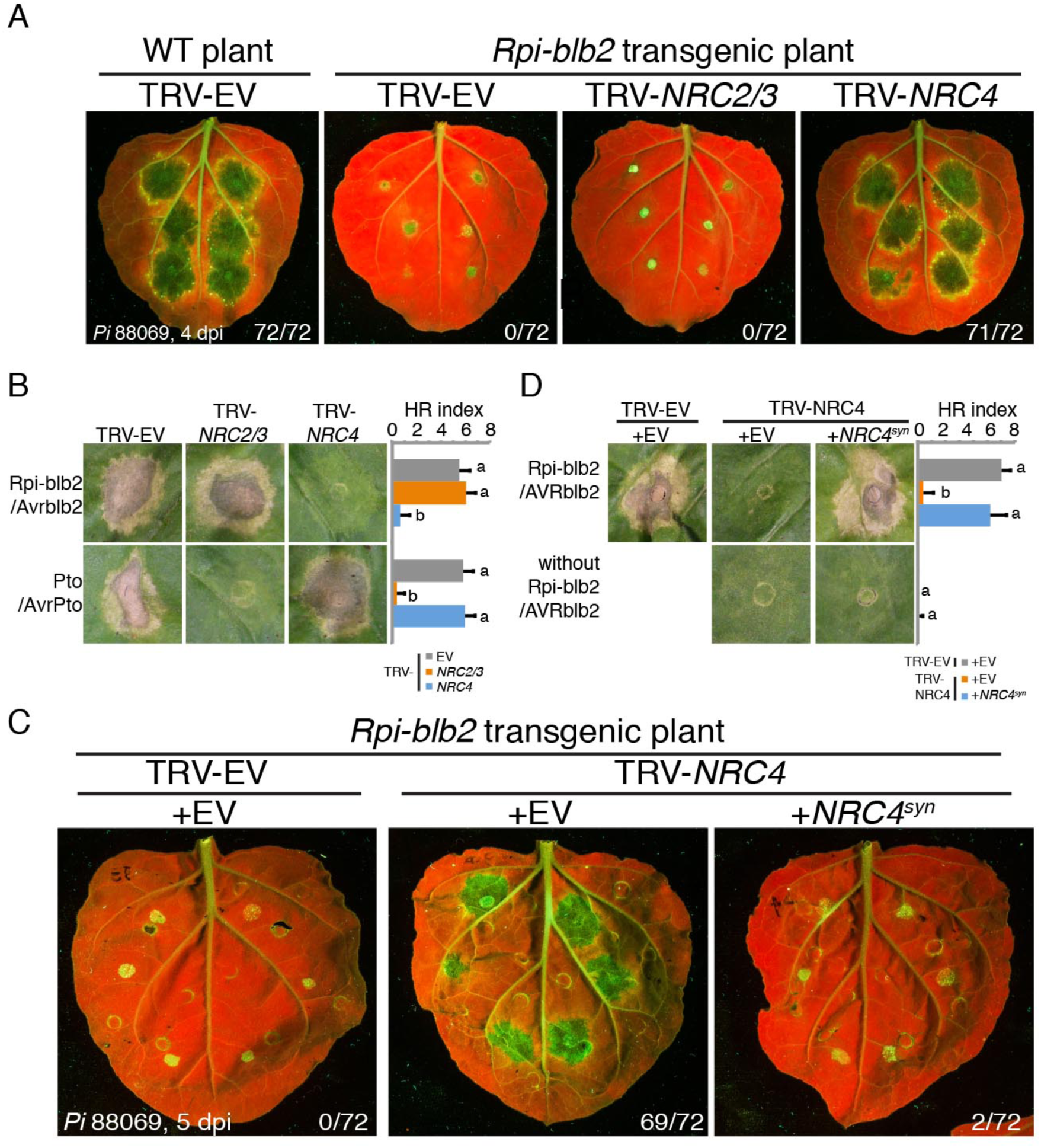
*NRC4* is required for Rpi-blb2-mediated immunity. (**A**) Silencing of *NRC4* compromises Rpi-blb2-mediated resistance. *Phytophthora infestans* strain 88069 (*Pi* 88069) was inoculated on *Rpi-blb2* transgenic *Nicotiana benthamiana* pre-infected with *Tobacco rattle virus* (TRV) to silence *NRC2/3* or *NRC4*. Wild type (WT) plant with TRV empty vector (TRV-EV) was used as a susceptible control. Experiments were repeated 3 times with 24 inoculation sites each time. The numbers on the right bottom indicate the sum of spreading lesions/total inoculation sites from the three replicates. Images were taken under UV light at 4 days post inoculation (dpi). (**B**) Silencing of *NRC4* compromises Rpi-blb2- but not Prf-mediated hypersensitive cell death. Rpi-blb2/AVRblb2 or Pto/AvrPto (cell death mediated by Prf) were co-expressed in *NRC2/3-* or *NRC4*-silenced plants by agroinfiltration. Hypersensitive response (HR) was scored at 7 days after agroinfiltration. Bars represent mean + SD of 24 infiltration sites. Statistical differences among the samples were analyzed with ANOVA and Tukey’s HSD test (p-value < 0.001). (**C**) Expression of silencing-resilient synthetic *NRC4* (*NRC4^syn^*) rescues Rpi-blb2-mediated resistance in *NRC4*-silenced plants. Experiments were repeated 3 times with 24 inoculation sites each time. The numbers on the right bottom indicate the sum of spreading lesion/total inoculation sites from the three replicates. Images were taken under UV light at 5 days post inoculation (dpi). (**D**) Expression of silencing-resilient synthetic *NRC4* (*NRC4^syn^*) rescues Rpi-blb2-mediated cell death in *NRC4*-silenced plants. Hypersensitive response (HR) was scored at 7 days after agroinfiltration. Bars represent mean + SD of 24 infiltrations sites. Statistical differences among the samples were analyzed with ANOVA and Tukey’s HSD test (p-value < 0.001).

NRC4 defines a distinct clade within the NRC family (Fig. S3A)(18). Of the 9 *NRC* genes in *N. benthamiana*, four were expressed to significant levels in leaves but only *NRC4* transcript levels were reduced in *NRC4*-silenced plants (Fig. S1D, Fig. S3B). Among the expressed genes, *NRC2* and *NRC3* are required for bacterial resistance mediated by the NLR protein Prf in *N. benthamiana* (9, 21) but were not essential for Rpi-blb2 functions in our silencing experiments (Fig. 1A-B). In contrast, *NRC4* was not essential for Prf-mediated cell death and resistance to the bacterial pathogen *Pseudomonas syringae* (Fig. 1B; Fig. S4).

Phylogenetic analyses of the complete repertoire of NLR proteins from the solanaceous plants tomato, potato, pepper and *N. benthamiana* revealed that the NRC family groups with the Rpiblb2 and Prf clades in a well-supported superclade (Fig. S5). Interestingly, this superclade includes additional well-known NLR, such as Rx (22, 23), Bs2 (24), R8 (25), Sw5b (26), R1 (27) and Mi-1.2 (28), which confer resistance to diverse plant pathogens and pests (Fig. S5; Table S1). This prompted us to test the extent to which NRC proteins are involved in immune responses mediated by these phylogenetically related disease resistance proteins.

Silencing of *NRC2* and *NRC3* only affected Prf and did not alter the hypersensitive cell death mediated by 13 other NLR proteins (Fig. 2). In contrast, silencing of *NRC4* compromised the hypersensitive cell death mediated by Mi-1.2 (28), an Rpi-blb2 ortholog that provides resistance to nematodes and insects; CNL-11990^D474V^ (18), an autoactive mutant of a CNL (NLR with a N-terminal coiled-coil domain) of unknown function, and R1 (27), an NLR that confers resistance to *P. infestans* (Fig. 2, Fig. S6A). Further, *NRC4* silencing abolished R1-mediated disease resistance to *P. infestans* and the phenotype was rescued by a silencing-resilient synthetic *NRC4* gene (Fig. S6B-D).

**Fig. 2.**
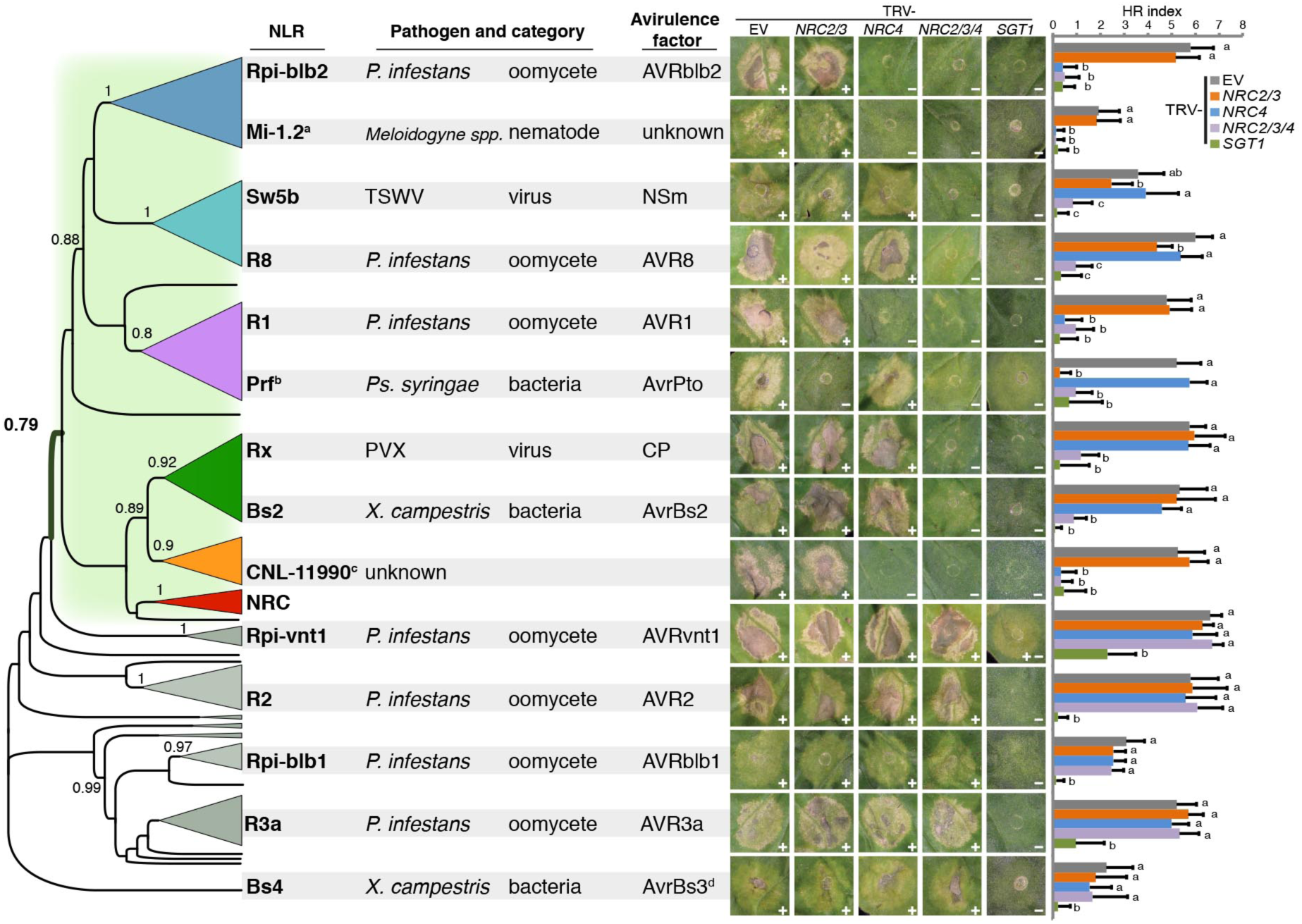
**NRC clade and its sister clades form a complex signaling network Left panel:** phylogenetic tree of NLR proteins identified from genomes of solanaceous plants, simplified from Fig. S5. **Middle panel**: list of pathogens and avirulence effectors (AVR) sensed by the corresponding NLR immune receptors. TSWV, tomato spotted wilt virus; *Ps*., *Pseudomonas*; PVX, *Potato virus X*; *X.*, *Xanthomonas.* **Right panel**: analysis of hypersensitive cell death mediated by different solanaceous NLR proteins in *NRC*-silenced plants. Different NLR and AVR effector combinations were expressed in control (EV), *NRC2/3*-, *NRC4*-, *NRC2/3/4*- and *SGT1*-silenced plants by agroinfiltration. “+” indicates cell death phenotype was observed. “−” indicates cell death phenotype was compromised. Hypersensitive response (HR) was scored at 7 days after agroinfiltration. Bars represent mean + SD of 24 infiltration sites. Statistical differences among the samples were analyzed with ANOVA and Tukey’s HSD test (P-value < 0.001). ^a^Pathogen proteins sensed by Mi-1.2 have not been identified yet. Hence, the autoactive mutant Mi-1.2^T557S^ was used here. ^b^Co-expression of Pto and AvrPto was used for testing Prf-mediated cell death. ^c^CNL-11990, a CNL cloned from tomato, has no assigned function. The autoactive mutant CNL-11990^D474V^ was used here. ^d^Bs4 senses both AvrBs3 and AvrBs4 from *X. campestris*. AvrBs3 was used here.

Given that the three expressed NRC proteins share extensive sequence similarity (Fig. S7), we hypothesized that NRC2, NRC3 and NRC4 are functionally redundant for additional NLR in the “NRC” superclade (Fig. 2). To test our hypothesis, we simultaneously silenced the three *NRC* genes and discovered that silencing compromised hypersensitive cell death mediated by Sw5b, R8, Rx and Bs2 in addition to the 5 NLR mentioned above (Fig. 2, Fig. S8, Fig. S9). In contrast, the triple *NRC* silencing did not affect hypersensitive cell death mediated by the 5 tested NLR that map outside the NRC superclade (Fig. 2) and did not abolish resistance to *P. infestans* conferred by two of these NLR proteins (Fig. S10).

We validated NRC2, NRC3 and NRC4 redundancy by complementation in the triple silencing background with silencing-resilient synthetic *NRC* (Fig. S11). This confirmed that the three NRC proteins display specificity to Rpi-blb2 and Prf but have redundant functions in Rx, Bs2, R8 and Sw5b mediated hypersensitive cell death (Fig. S11).

To further validate that NRC2, NRC3 and NRC4 redundantly contribute to immunity, we examined the resistance mediated by Rx to *Potato virus X* (PVX) (22, 23) in plants silenced for single, double or triple combinations of *NRC* genes (Fig. S12). Rx-mediated resistance to PVX was only abolished in the triple silencing background resulting in systemic spread and accumulation of the virus (Fig. S12, Fig. S13). Remarkably, silencing-resilient synthetic *NRC2*, *NRC3* and *NRC4* individually complemented the loss of resistance to PVX in triple *NRC*-silenced plants confirming their functional redundancy (Fig. S14). This and previous results indicate that the three NRC proteins display varying degrees of redundancy and specificity towards the 9 NLR revealing a complex immune signaling network (Fig. S15).

Our observation that NRC proteins and their NLR mates are phylogenetically related (Fig. S5) prompted us to reconstruct the evolutionary history of the NRC superclade. Higher order phylogenetic analyses of complete NLR repertoires from representative plant taxa revealed that the NRC superclade is missing in rosids but present in all examined caryophyllales (sugar beet) and asterids (kiwifruit, coffee, monkey flower, ash tree and Solanaceae species) (Fig. S16, Fig. 3A-B, Fig. S17-20). Interestingly, sugar beet and kiwifruit, the early branching species, have only a single protein that groups with the NRC family, along with 2 and 4 NLR that cluster with the NRC-dependent NLR (Fig. 3A-B, Fig. S20). The dramatic expansion of the NRC superclade started prior to the divergence of Gentianales (coffee) from other asterids about 110-100 million years ago (29, 30) to account for over one half of all NLR in some of the species (Fig. 3B). We conclude that the NRC superclade evolved from an ancestral pair of genetically linked NLR genes, as in sugar beet, to duplicate and expand throughout the genomes of asterid species into a complex genetic network that confers immunity to a diversity of plant pathogens (Fig. 3C-D, Fig. S21).

**Fig. 3.**
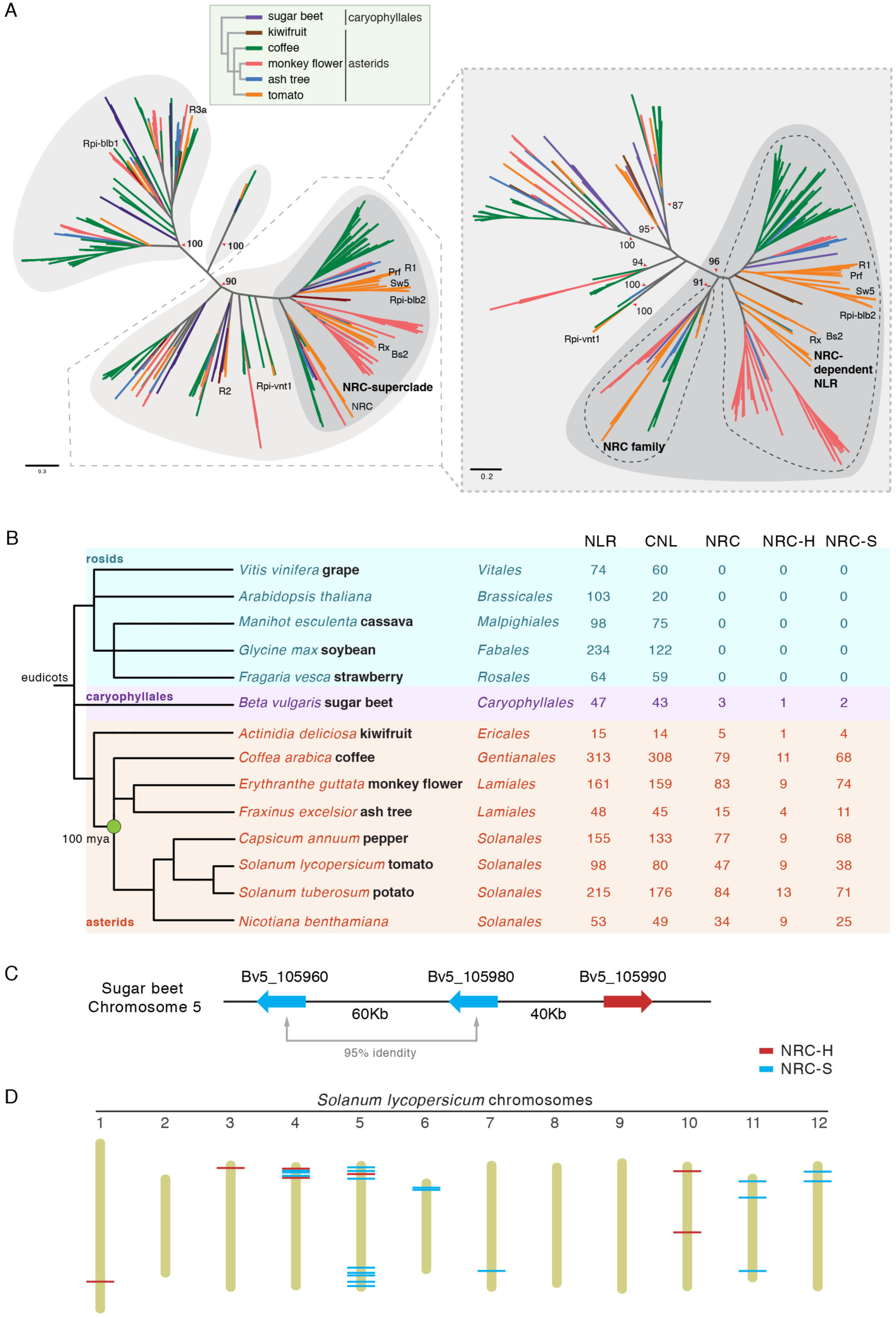
NRC-superclade emerged from a NLR pair over 100 million years ago. (**A**) Phylogeny of CNL (CC-NLR) identified from asterids (kiwifruit, coffee, monkey flower, ash tree and tomato) and caryophyllales (sugar beet). Sequences identified from different species are marked with different color as indicated. The bootstrap supports of the major nodes are shown. The phylogenetic tree in the right panel, which includes only sequences from the indicated lineages in the left panel, shows that the NRC sequences form a well-supported superclade that occurs in asterids and caryophyllales. The scale bars indicate the evolutionary distance in amino acid substitution per site. Details of the full phylogenetic tree can be found in Fig. S19-20. (**B**) Summary of phylogeny and number of NLR identified in the different plant species. Phylogenetic tree of plant species was generated by using phyloT based on National center for Biotechnology Information (NCBI) taxon identification numbers. Numbers of NLR identified in each category were based on NLR-parser and the phylogenetic trees in (**A**) and Fig. S16-20. Mya, million years ago; CNL, CC-NLR; NRC, NRC-superclade; NRC-H, NRC family (helper NLR); NRC-S, NRC-dependent NLR (sensor NLR). (**C**) Schematic representation of the NRC gene cluster on sugar beet chromosome 5. The two NRC-S paralogs are marked in blue, and the NRC-H gene is marked in red. (**D**) Physical map of NRC superclade genes on tomato chromosomes. The NRC-S paralogs are marked in blue, and the NRC-H paralogs are marked in red. The detail information of the physical map can be found in Fig. S21.

Thus, NLR pairs can evolve into a signalling network with a complex architecture. However, NLR evolution must be constrained by their mode of action. The selective pressures shaping the evolution of NLR pairs that operate by negative regulation would limit their expansion due to the genetic load caused by autoimmunity (Fig. S22). The NRC family appears to function through a mechanism that accommodates evolutionary plasticity beyond genetically linked pairs of NLR.

Genetic redundancy increases robustness of signaling networks (31-33). The NRC network may therefore augment the plant capacity to counteract rapidly evolving pathogens. Multiple NRC would further enhance evolvability of sensor NLR, i.e. their capacity to undergo rapid adaptive evolution. Harnessing the processes that underpin NLR network structure and function would open up new approaches for developing disease resistant crops.

## Acknowledgments

We thank Oliver Furzer, Jonathan Jones, John Rathjen, Brian Staskawicz, Geert Smant, Sebastian Schornack, Frank Takken, Vivianne Vleeshouwers and Cyril Zipfel for providing materials and technical supports. We are grateful to Lida Derevnina, Yasin Dagdas, Benjamin Petre, Erin Zess and Esther van der Knaap for helpful suggestions. This project was funded by the Gatsby Charitable Foundation, Biotechnology and Biological Sciences Research Council (BBSRC), and European Research Council (ERC).

## References

1. J. L. Dangl, D. M. Horvath, B. J. Staskawicz, Pivoting the plant immune system from dissection to deployment. Science 341, 746–751 (2013).

2. J. von Moltke, J. S. Ayres, E. M. Kofoed, J. Chavarria-Smith, R. E. Vance, Recognition of bacteria by inflammasomes. Annual Review of Immunology 31, 73–106 (2013).

3. P. N. Dodds, J. P. Rathjen, Plant immunity: towards an integrated view of plant-pathogen interactions. Nat Rev Genet 11, 539–548 (2010).

4. F. Jacob, S. Vernaldi, T. Maekawa, Evolution and conservation of plant NLR functions. Front Immunology 4, 297 (2013).

5. R. M. Clark et al., Common sequence polymorphisms shaping genetic diversity in *Arabidopsis thaliana*. Science 317, 338–342 (2007).

6. L. McHale, X. P. Tan, P. Koehl, R. W. Michelmore, Plant NBS-LRR proteins: adaptable guards. Genome Biol 7, 212 (2006).

7. J. Win et al., Effector biology of plant-associated organisms: concepts and perspectives. Cold Spring Harb Symp Quant Biol 77, 235–247 (2012).

8. V. Bonardi et al., Expanded functions for a family of plant intracellular immune receptors beyond specific recognition of pathogen effectors. Proc Natl Acad Sci USA 108, 16463–16468 (2011).

9. C. H. Wu, K. Belhaj, T. O. Bozkurt, M. S. Birk, S. Kamoun, Helper NLR proteins NRC2a/b and NRC3 but not NRC1 are required for Pto-mediated cell death and resistance in *Nicotiana benthamiana*. New Phytol 209, 1344–1352 (2016).

10. V. Bonardi, K. Cherkis, M. T. Nishimura, J. L. Dangl, A new eye on NLR proteins: focused on clarity or diffused by complexity? Curr Opin Immunol 24, 41–50 (2012).

11. T. K. Eitas, J. L. Dangl, NB-LRR proteins: pairs, pieces, perception, partners, and pathways. Curr Opin Plant Biol 13, 472–477 (2010).

12. S. Cesari, M. Bernoux, P. Moncuquet, T. Kroj, P. N. Dodds, A novel conserved mechanism for plant NLR protein pairs: the "integrated decoy" hypothesis. Front Plant Sci 5, 606 (2014).

13. T. Sarkinen, L. Bohs, R. G. Olmstead, S. Knapp, A phylogenetic framework for evolutionary study of the nightshades (Solanaceae): a dated 1000-tip tree. BMC Evol Biol 13, 214 (2013).

14. G. van Ooijen, H. A. van den Burg, B. J. C. Cornelissen, F. L. W. Takken, Structure and function of resistance proteins in solanaceous plants. Annu Rev Phytopathol 45, 43–72 (2007).

15. V. G. A. A. Vleeshouwers et al., Understanding and exploiting late blight resistance in the age of effectors. Annual Review of Phytopathology, 49, 507–531 (2011).

16. E. A. G. van der Vossen et al., The *Rpi-blb2* gene from *Solanum bulbocastanum* is an *Mi-1* gene homolog conferring broad-spectrum late blight resistance in potato. Plant J 44, 208–222 (2005).

17. S. K. Oh et al., In planta expression screens of *Phytophthora infestans* RXLR effectors reveal diverse phenotypes, including activation of the *Solanum bulbocastanum* disease resistance protein Rpi-blb2. Plant Cell 21, 2928–2947 (2009).

18. Materials and methods are available as supplementary material.

19. S. J. Williams et al., Structural basis for assembly and function of a heterodimeric plant immune receptor. Science 344, 299–303 (2014).

20. S. Cesari et al., The NB-LRR proteins RGA4 and RGA5 interact functionally and physically to confer disease resistance. EMBO J 33, 1941–1959 (2014).

21. A. Balmuth, J. P. Rathjen, Genetic and molecular requirements for function of the Pto/Prf effector recognition complex in tomato and *Nicotiana benthamiana*. Plant J 51, 978–990 (2007).

22. A. Bendahmane, K. Kanyuka, D. C. Baulcombe, The *Rx* gene from potato controls separate virus resistance and cell death responses. Plant Cell 11, 781–791 (1999).

23. W. I. L. Tameling, D. C. Baulcombe, Physical association of the NB-LRR resistance protein Rx with a ran GTPase-activating protein is required for extreme resistance to *Potato virus X*. Plant Cell 19, 1682–1694 (2007).

24. T. H. Tai et al., Expression of the *Bs2* pepper gene confers resistance to bacterial spot disease in tomato. Proc Natl Acad Sci USA 96, 14153–14158 (1999).

25. J. H. Vossen et al., The *Solanum demissum R8* late blight resistance gene is an *Sw-5* homologue that has been deployed worldwide in late blight resistant varieties. Theor Appl Genet 129, 1785–1796 (2016).

26. S. H. Brommonschenkel, A. Frary, A. Frary, S. D. Tanksley, The broad-spectrum tospovirus resistance gene *Sw-5* of tomato is a homolog of the root-knot nematode resistance gene *Mi*. Mol Plant Microbe Interact 13, 1130–1138 (2000).

27. A. Ballvora et al., The *R1* gene for potato resistance to late blight (*Phytophthora infestans*) belongs to the leucine zipper/NBS/LRR class of plant resistance genes. Plant J 30, 361–371 (2002).

28. S. B. Milligan et al., The root knot nematode resistance gene *Mi* from tomato is a member of the leucine zipper, nucleotide binding, leucine-rich repeat family of plant genes. Plant Cell 10, 1307–1319 (1998).

29. K. Bremer, E. M. Friis, B. Bremer, Molecular phylogenetic dating of asterid flowering plants shows early Cretaceous diversification. Syst Biol 53, 496–505 (2004).

30. N. Wikstrom, K. Kainulainen, S. G. Razafimandimbison, J. E. E. Smedmark, B. Bremer, A revised time tree of the asterids: establishing a temporal framework for evolutionary studies of the coffee family (Rubiaceae). Plos One 10, e0126690 (2015).

31. A. Wagner, Robustness and evolvability in living systems. Princeton studies in complexity (Princeton University Press, Princeton, N.J., 2005), 367 p.

32. M. A. Fares, The origins of mutational robustness. Trends Genet 31, 373–381 (2015).

33. H. Kitano, Biological robustness. Nat Rev Genet 5, 826–837 (2004).

